# Asporin Improves Cardiac Myocyte Response to Ischemia and Reperfusion Stress

**DOI:** 10.64898/2026.01.22.701197

**Authors:** Deepika Rai, Mukta Basu, Liam McCarthy, Divya Gupta, Ashley Dinh, Matthew Aryes, Ajay Bhardwaj, Pavithra Nedumaran, Reetu Thakur, Aleksandar B Stotland, Honit Piplani, Sarah J Parker

## Abstract

**Background:** Myocardial Infarction (MI) remains a leading cause of mortality worldwide, despite advancements in clinical therapies and interventions. MI results from prolonged ischemia, leading to hypoxia-induced damage to cardiac tissue and reperfusion-injury (R/I) that aggravates cardiomyocyte (CM) loss. One key cellular event during this process is accumulation of dysfunctional mitochondria, resulting from environmental hypoxia and subsequent oxidative stress upon reperfusion. Post-MI cardiac-remodeling involves changes in both cellular and extracellular matrix (ECM). Ubiquitin-dependent and independent autophagy are crucial for cardio protection during this phase. The ECM provides structural integrity and functions as a reservoir for signaling molecules. Asporin (ASPN), a small leucine-rich proteoglycan, plays a role in modulating cardiac-remodeling by limiting excessive fibrosis and protecting CMs from cell death.

**Methods:** We investigated the therapeutic potential of ASPN by using an exogenous recombinant peptide of ASPN (rASPN), testing its effects using an *in-vitro* ischemia-reperfusion (I/R) model simulating MI conditions. Two I/R models were developed using an immortalized human embryonic cardiac cell line to reflect the hypoxia-reperfusion (H/R) phases of MI. In the No-Reoxygenation (No-ReOx) model, cells were subjected to hypoxia for 18 hours, with or without exogenous rASPN. In the Reoxygenation (ReOx) model, cells underwent 18 hours of hypoxia, then 12 hours of reoxygenation (simulating reperfusion), with or without rASPN.

**Results:** Proteomics revealed that ASPN modulates key pathways involved in apoptosis, non-canonical autophagy, and metabolic reprogramming. Additionally, ASPN influenced immune response pathways and significantly affected TGF-β signaling, a central mediator of cardiac fibrosis and remodeling post-MI. These findings indicate that ASPN plays a multifaceted role in regulating cellular responses to hypoxia and R/I.

**Conclusions:** Our H/R model simulates key aspects of MI and R/I. The protective role of ASPN observed in this model suggests it as a promising candidate for developing cardioprotective therapies to minimize R/I and adverse cardiac-remodeling following MI.

## Introduction

Myocardial infarction (MI), primarily resulting from the rupture of atherosclerotic plaques and subsequent thrombotic occlusion of coronary arteries, remains one of the leading causes of mortality worldwide [1, 2]. The ensuing ischemia leads to myocardial hypoxia, which is a major cellular stressor that initiates cardiomyocyte (CM) death through apoptosis and necrosis [3-5]. Timely reperfusion is the standard clinical intervention for restoring blood flow and minimizing infarct size [6, 7]. Paradoxically, however, the sudden influx of oxygenated blood can cause additional cellular injury, known as ischemia-reperfusion (I/R) injury [8]. This reperfusion-associated oxidative stress contributes to further CM damage, limits myocardial recovery, and worsens clinical outcomes [9].

Understanding the molecular basis of I/R injury is crucial for developing more effective therapeutic strategies [10]. One of the primary targets of hypoxic and reperfusion-induced damage is the mitochondrion, a vital organelle responsible for ATP production and maintaining energy homeostasis [11]. CMs, with their high metabolic demand, are particularly vulnerable to mitochondrial dysfunction[11, 12]. Hypoxia and I/R stress lead to oxidative damage to mitochondrial DNA and impaired mitochondrial function, resulting in an energy imbalance, increased reactive oxygen species (ROS), and activation of cell death pathways [13]. The accumulation of dysfunctional mitochondria disrupts cellular homeostasis and contributes to adverse cardiac remodeling after myocardial infarction (MI) [13].

Autophagy, particularly mitophagy, serves as a critical quality control mechanism by removing damaged mitochondria and recycling cellular components[13]. Emerging evidence suggests that enhancing autophagy can attenuate CM apoptosis and improve functional recovery following MI, leading to the hypothesis that targeting autophagic flux during I/R is a promising target for cardioprotective interventions [14].

Our previous work has identified the extracellular matrix (ECM) protein Asporin (ASPN) as a novel regulator of autophagy in cardiac fibroblasts[15]. Genetic deletion of ASPN in mice led to increased infarct size, exacerbated fibrosis, elevated CM death, and impaired cardiac function after permanent coronary artery ligation (PCAL)[15]. *In vitro* studies in cultured fibroblasts demonstrated that ASPN interacts with the TGF-β1 signaling pathway—a central mediator of fibrosis acting as a negative feedback regulator by suppressing SMAD2/3 phosphorylation[15]. While TGFβ1 is initially protective post-injury, sustained activation promotes fibrosis and maladaptive remodeling[15]. These findings position ASPN as a dual-function modulator of both autophagy and fibrotic signaling in cardiac tissue[15]. While these studies focused on fibroblasts are enlightening, it remains unclear whether ASPN exerts similar regulatory roles in human CMs, particularly in the context of MI-related hypoxia and reperfusion. Furthermore, the broader proteomic effects and potential therapeutic applications of recombinant ASPN (rASPN) in CMs have not been fully elucidated.

In this work, we examine the molecular and functional impact of exogenous rASPN on human CMs under simulated hypoxic and reperfusion conditions. Using clinically relevant human cell *in vitro* modeling and a murine PCAL model, we explore the role of rASPN in modulating autophagy, apoptosis, and TGF-β signaling. Through deep proteomic profiling (LC-MS/MS) and integrative bioinformatics analysis, we uncover novel mechanistic insights into the cardioprotective effects of rASPN, highlighting its therapeutic potential for mitigating ischemia-reperfusion (I/R) injury and improving cardiac outcomes post-myocardial infarction (MI).

## Experimental procedures

### Cell Culture

AC16 human ventricular CMs cells were obtained from ATCC (CRL-3568) and maintained as described previously in growth media Dulbecco’s modified Eagle’s medium (DMEM/F-12 (1:1)) supplemented with 10% fetal bovine serum (FBS), antibiotics, and antimycotics, pH 7.4 at 37°C in the absence of 5% CO_2_ [16]. The hypoxia/reoxygenation (H/R) model was established to simulate I/R injury *in vitro*, wherein confluent cells (10mm culture dishes) were transferred to a hypoxic modular chamber (0.1% oxygen) for 18 h at 37 °C. After hypoxia, the medium was replaced with a fresh medium, and the dishes were transferred to a normoxic incubator (∼21% oxygen) for further assessment of the effects of 12 hours of reoxygenation. For recombinant asporin peptide (rASPN), cell culture media were supplemented with peptide at a concentration of 50 ng/mL either during hypoxia exposure or after hypoxia during the 12-hour recovery phase. In the case of chloroquine (Sigma Aldrich-C1650000) treatment, cells were subjected to 50µM concentrations, and recovery monitoring was extended to 24 hours after hypoxia. A diagram of the overall study design is provided in Figure 1A.

**Figure 1.**
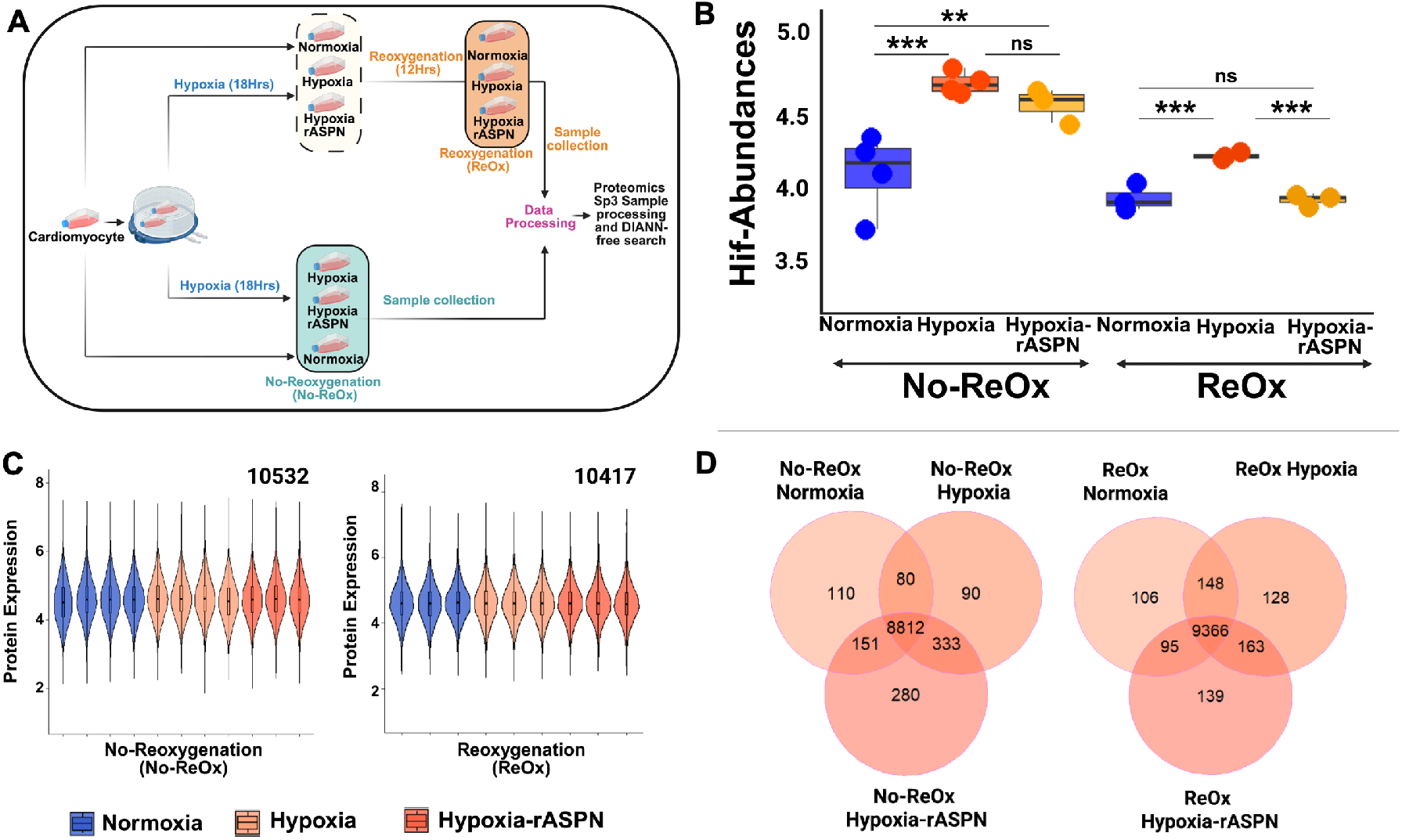
Summary of experimental design and protein expression analysis through bulk-proteomics in AC-16 CMs under hypoxia and treatment conditions. (A) Schematic illustration of the experimental design; AC-16 CMs were subjected to two ischemic models: Ischemia without reoxygenation (No-ReOx) and Ischemia followed by reoxygenation (ReOx). (B) Abundance of HIF-1α protein across experimental conditions (C) Violin plots of intensity distrubutions for all proteins detected across for Normoxia, Hypoxia, and Hypoxia-rASPN in both the No-ReOx and ReOx experiments. (D) Venn diagram for overlapping and uniquely identified proteins across experimental groups in both models, No-ReOx and ReOx. rASPN = recombinant asporin peptide. No-ReOx = no reoxygenation after acute hypoxia; ReOx = 12-hours of reoxygenation exposure after hypoxia. HIF1 = Hypoxia Inducible Factor 1.

### Mouse model of permanent coronary ligation to assess the impact of ASPN on WT and ASPN-KO mice

*In vivo* tissue data were analyzed using the hearts of mice collected according to the methodologies described in our previous study [15], with full details provided in that publication. Briefly, a mouse model was used to assess the effect of ASPN KO on mice after permanent coronary artery ligation on the development of LV remodeling. Mice were anesthetized with ketamine (50 mg/kg)/xylazine (10 mg/kg), and anesthesia was maintained using 0.5% isoflurane mixed with 1.0 L/min 100% O_2_ by mechanical ventilation. A left thoracotomy was performed to expose the heart, followed by the opening of the pericardium. Then, a 7–0 silk suture was placed around the proximal left coronary artery 2 mm below the left atrial appendage using a curved 27.5 G needle. The vessel was ligated, the chest closed, and the animals allowed to recover for 24 hours. Control mice were subjected to all procedures, including suture placement, with the only difference being that no actual ligation was performed (Sham).

### Western blotting

Protein concentration was estimated using a Thermofisher-Pierce bicinchoninic acid (BCA) Protein Assay kit. The identical protein concentration from each condition compared was resolved on Bolt 4-12% Bis-Tris Plus gels (Bio-Rad Laboratories Inc.) and transferred to nitrocellulose membranes. Total protein loading was assessed using Ponceau S stains [15]. Membranes were blocked with 2.5% BSA in Tris-Buffered Saline with Tween 20 (TBST) for an hour, then incubated with primary antibody (1:1000 diluted) against HIF1-α (Catalog # EP1215Y, AbCam), BNIP3 (Catalog # EP1215Y, Cell Signaling), LC3-I/II (EP1215Y, Cell Signaling), BAX (Catalog # 2772, Cell Signaling), BCL2 (Catalog # D55G8,4223, Cell Signaling), SMAD-2/3 (Catalog # ab217553, AbCam), pSMAD-2/3 (Catalog # 8828, Cell Signaling), or β-Actin (Catalog # 4970, Cell Signaling) at 4ºC overnight. Furthermore, the membranes were washed with TBST at room temperature and incubated with KPL peroxidase-labeled secondary antibodies for 2 hours at the same temperature. Washing thrice with TBST and developing immunoreactive bands with clarity western ECL substrate (Bio-Rad Laboratories Inc.). Images were captured using a ChemiDoc XRS system (Bio-Rad Laboratories Inc.) and analyzed using NIH ImageJ software for densitometry.

### Immunofluorescence

Cells were first fixed with 4% paraformaldehyde (PFA) in phosphate-buffered saline (PBS) for 20 min and washed twice with PBS. Permeabilization and blocking of tissue sections or fixed cells were performed for 1 h in a “blocking buffer” containing PBS with 10% donkey serum (Millipore) and 0.1% Triton-X (Bio-Rad) [17]. Primary antibodies were diluted in the blocking buffer and kept on the cells overnight at 4°C. The following primary antibodies and dilutions were used: LC-3 (1:2000, Abcam), Cleaved Caspase 3 (1:500, Thermofisher Scientific), Troponin T (TnT) (1:200, Thermofisher Scientific), and Smad4 (1:200, Abcam). The next day, after thorough washing with PBS containing 0.1% Tween-20 (Thermo Fisher), cells were incubated with species-specific Alexa Fluor-conjugated secondary antibodies (Thermo Fisher) diluted in blocking buffer (1:1000) for 1 hour at room temperature. After washing in PBS with 0.1% Tween-20, cells were incubated with DAPI diluted in PBS (1:2500) for 15 min. Immunofluorescence images were taken using appropriate fluorescent filters using the Revolve-Echo-M-00151 Microscope (Magnification was 40 µm; A total of seven replicates for each condition have been used, and a total of 20 images per condition have been analyzed). For image analysis, we used Python with scikit-image, NumPy, Matplotlib, and Pandas libraries. Further RGB images were processed using Python by extracting the green channel (LC-3) and the red channel (CC-3) and then segmenting the foreground signals using Otsu thresholding. Followed by an assessment of connected components using an adaptive size criterion (10 pixels minimum; maximum = mean area + 2 SD) to remove noise and non-specific objects. Valid puncta were counted per channel, visually validated with overlays, and quantitative results were exported to Excel for statistical analysis.

### Immunohistochemistry (fluorescence)

As noted above, mouse tissue from our previous publication was used for *in vivo* confirmation. Details regarding the animal model and genetics can be found in [15]. For our studies, formalin-fixed paraffin-embedded cross-sections of hearts from wild-type and ASPN knock-out (KO) mice with or without permanent coronary artery ligation (PCAL). Slide sections were deparaffinized with xylene treatment twice and ethanol gradient from 100% to 70%, followed by hydration with deionized water. Followed by hydration in deionized water. Sections were placed in a container with antigen retrieval solution (Universal HIER antigen retrieval reagent (10X), AbCam) at 95°C for 30 min, followed by 30 min at RT [15, 18]. Permeabilization and blocking were done using protein blocking solution (Protein Block, AbCam). The following primary antibodies and dilutions were used: LC-3 (1:200, Abcam), Cleaved Caspase 3 (1:200, Thermofisher Scientific), Troponin T (TnT) (1:200, Thermofisher Scientific), and Smad4 (1:200, Abcam), all of which were incubated overnight at 4 °C. The next day, after thorough washing with PBS containing 0.1% Tween-20 (Thermo Fisher), cells were incubated with species-specific Alexa Fluor-conjugated secondary antibodies (Thermo Fisher) diluted in blocking buffer (1:1000) for 1 hour at room temperature. After washing in PBS with 0.1% Tween-20, mounting with DAPI (Vector shield, Vector Laboratories). Immunofluorescence images were taken using appropriate fluorescent filters using the Revolve-Echo-M-00151 Microscope (Magnification was 40 µm; A total of three heart sample replicates for each condition have been used, and a total of 20 images per condition have been analyzed) [19, 20]. For image analysis, we utilized Python, incorporating scikit-image, SciPy, NumPy, Matplotlib, and Pandas. Further RGB images were processed using Python by extracting the green channel (LC-3) and the red channel (CC-3) and then segmenting the foreground signals using Otsu thresholding. Calculations were performed in the same workflow as for cells stained with LC-3 and CC-3, as mentioned above. In case of SMAD-4 puncta calculations, we processed all three channel RGB in Python by extracting the green channel (LC-3) and the red channel (CC-3), with nuclei segmented from the DAPI channel using Otsu thresholding and morphological cleanup. CMs and respective nuclei were identified based on the characteristic nuclear size and elongation, expanded by dilation to approximate cell territories, and refined using spatial proximity to the red-channel signal to improve specificity. Red punctas (CT-T stained) were enhanced, segmented, and separated using watershed analysis, with small objects excluded. Each punctum was classified according to its spatial relationship with cardiomyocyte masks, nuclear regions, and red-channel signal into cardiomyocyte nuclear, cardiomyocyte cytoplasmic, cardiomyocyte-only, or non-cardiomyocyte categories. Segmentation and classification results were visually validated, and per-punctum measurements and quantitative summaries were exported to Excel for downstream analysis.

### Proteomic sample preparation and Mass spectrometry acquisition

As mentioned in the above section, total protein concentration was assayed, and 60 µg of protein per sample was aliquoted for proteomics sample processing. The Sp3 protocol[21] is used for protein processing, digestion, and cleaning, followed by drying samples by diluting them to 0.2 µg/µL for a total of 2 µg on the column, and then injecting them into a mass spectrometry (MS) system for analysis through LC-MS. DIA analysis was performed on an Orbitrap Astral (Thermo Scientific) mass spectrometer interfaced with an EASY-Spray™ nano-electrospray ionization source (Thermo Scientific, ES081) coupled to Vanquish Neo ultra-high-pressure chromatography system with 0.1% formic acid in water as mobile phase A and 0.1% formic acid in acetonitrile as mobile phase B. Peptides were separated at an initial flow rate of 1.2 µL/minute and a linear gradient of 4-9% B for 0-2 minutes, 9-25% B for 2-13 minutes, 25-35% B for 13-17 minutes. The flow rate was then increased to 3 µL/min, and the column was flushed with 35-99% B for 0.4 minutes, then held at 99% B for 0.5 minutes, and decreased to 5%. The column used was PepSep C18 15cm x 150 µm, 1.5µm (Bruker, P/N: 1893474). Source parameters were set to a voltage of 2300 V and a capillary temperature of 280°C. MS1 scan range was set to 360-1200 m/z, and MS1 resolution was set to 240,000 with an AGC target set to ‘Custom’ and a normalized AGC target set to 500%. The RF Lens was set to 40% with a maximum injection time of 10ms. The precursor mass range was set to 380-980 m/z, with 149 non-overlapping data-independent acquisition precursor windows of 4 m/z in size. MS2 scan range was set to 150-2000 m/z, and normalized HCD collision energy was set to 30%. Maximum injection time was set to 4ms with AGC target set to ‘Custom’ and normalized AGC target set to 200%. All data is acquired in profile mode using positive polarity.

### Proteomic analysis

Raw files obtained from the MS instrument computer were converted to mzML files using MSConvert software with peak picking enabled [22]. Data were analyzed using the DIA-NN software (version 1.8.1) with an in silico digested human-reviewed and canonical FASTA library downloaded from the UniProt database (December 2020). RT-dependent cross-run normalization was enabled[23]. Library generation was set to “FASTA digest for library-free search” and “Deep learning-based spectra, RTs and IMs prediction,” and MBR was turned on. When reporting protein numbers and quantities, the Protein.Group column in DIA-NN’s main report was used to identify the protein group and the Precursor.Normalized column was used to obtain the normalized precursor quantities. Next, the “Protein.Group” column was used to include only unique proteins that are quantified using proteotypic peptides. The precursor m/z range was set between 400 and 1000, and the fragment ion m/z range was set between 200 and 1800. Additionally, the missed cleavages rate was set to 2, and the peptide length range was set between 5 and 30. The protein inference parameter was set to “Protein Names”. All other settings were left at their default values. The software output was filtered at precursor q-value <1% and the global protein q-value <1% filter was also applied to all benchmarks. The precursor-level dataset was further filtered with a group-specific observation filter; precursors not meeting at least 50% of the samples are dropped. Protein and peptide-level quantification were then performed using the MSStats algorithm (v3.14), and protein-level comparison tests were performed using the MetaboanalystR package (v4.0.0).

### Statistical analysis

All statistical analysis and figure plots are performed using R (vMariposa Oersion 2025.05.1, mariposa orchid) and Python (version 4.26.1, Jupyter Notebook). The standard Student’s *t*-test was used to compare data from 2 groups only. Data with 3 or more groups was analyzed using Analysis of Variance (ANOVA) with Tukey post hoc test.

## Results

### Effect of ASPN on the Proteomic response of AC16-CMs to hypoxia and hypoxia-reperfusion

Our goal was to generate a more comprehensive understanding of how ASPN impacts human CMs during hypoxia and I/R. To achieve this, we utilized the established AC-16 cell line (adult human ventricular CMs) exposed to two ischemic experimental models, in the presence or absence of rASPN: (a) Ischemia without reoxygenation (no-reoxygenation), and (b) Ischemia followed by reperfusion (reoxygenation). Bulk proteomics was used to perform unbiased molecular profiling of the effects of recombinant rASPN on human AC16 CMs *in vitro* (Figure 1A).

Mass spectrometry (Figure 1B) analysis confirmed a significant effect of hypoxia in elevating hypoxia inducible factor 1-alpha (HIF1-α) protein levels, which remain significantly elevated throughout the ‘reperfusion’ recovery interval. Interestingly, rASPN blunted HIF1a levels, and that effect was most pronounced during the recovery phase (Figure 1B). Proteomic analysis was subsequently performed to quantify changes in protein abundance across conditions. As a quality control for sample loading on LC-MS and data normalization, the distribution of protein intensities was balanced across all sample sets as uniquely observed datasets (Figure 1C). A total of 10523 and 10417 proteins were observed in the hypoxia and recovery datasets, respectively, with substantial overlap in the proteins observed between conditions (Figure 1D). Under acute hypoxia, there were more proteins uniquely observed in the rASPN-treated cells relative to normoxia or hypoxia; however, after recovery, similar patterns of proteins were identified in only one condition (Figure 1D).

### Effect of rASPN on Cardiomyocyte Proteome during acute hypoxia

Differential abundance testing was performed on proteins shared across all conditions using pairwise comparisons between treatment groups, and significance determined using multiple testing corrected false discovery rate (FDR) < 0.05. In the No-ReOx condition, a total of 8812 (Figure 1C) proteins were identified across normoxia, hypoxia, and hypoxia-rASPN groups. Pairwise analysis between conditions revealed 1143 proteins with significant differential abundance (DA) in H vs N (14% of the detected proteome, Figure 2A). When compared to the non-hypoxic control, the HA condition also demonstrated a similar overall differential response (1181 DA proteins, 13% of the observed proteome, Figure 2B), reflecting the combined effect of both hypoxia and rASPN peptide. Between the HA vs. H, 424 DA proteins were observed (∼5% of the total detected proteome, Figure 2C), indicating the addition of rASPN in the presence of hypoxia alters the overall CM proteome beyond the effects of hypoxia alone.

**Figure 2.**
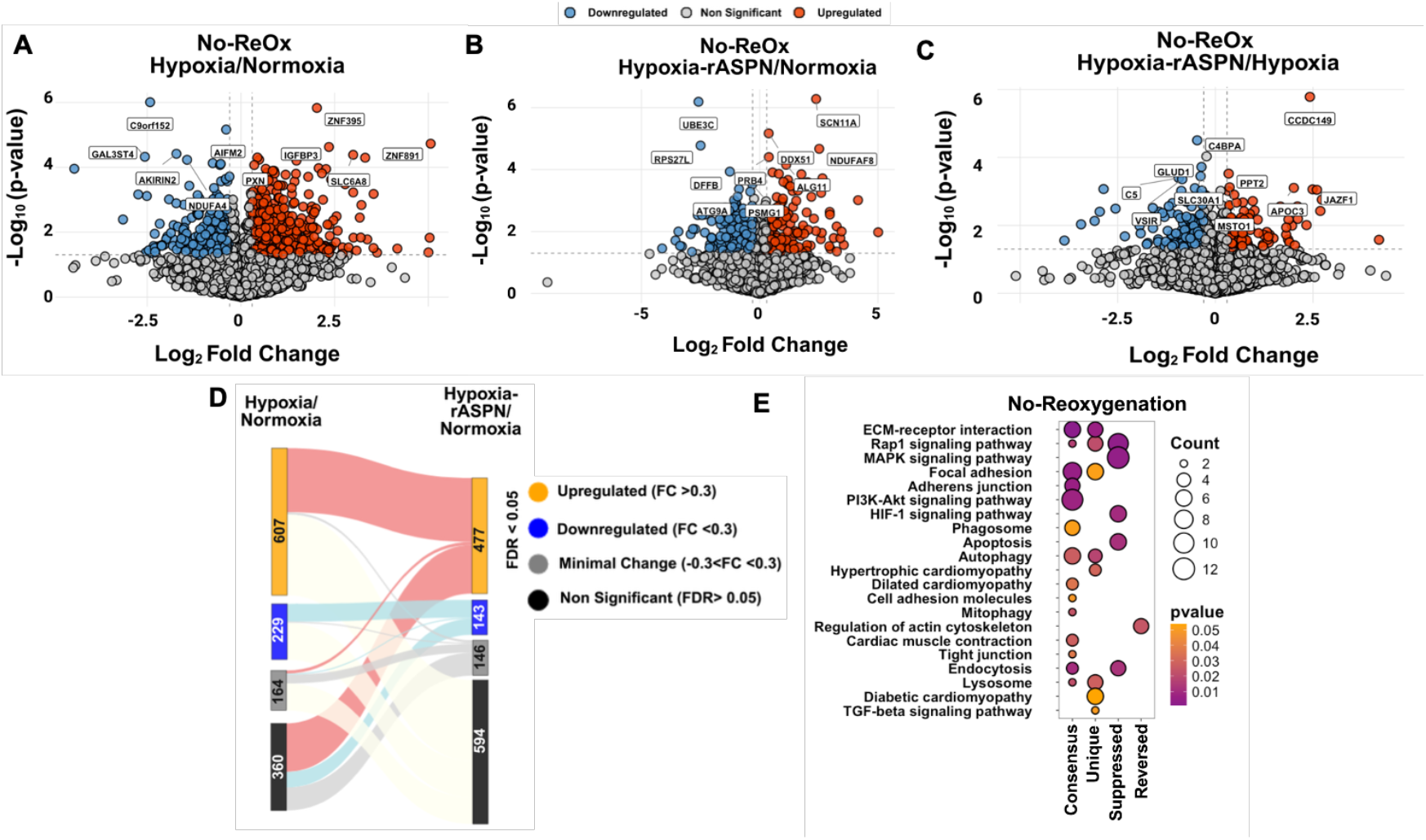
Overview of differentially expressed proteins (DEPs) identified in hypoxia and rASPN-treated conditions under No-ReOx. Volcano plot showing overall patterns of differential expression based on (A) the ratio of hypoxia to normoxia for No-Reoxygenation, (B) the ratio of hypoxia-rASPN to normoxia for No-Reoxygenation, and (C) the ratio of hypoxia-rASPN to hypoxia for No-Reoxygenation. For all volano plots, the x-axis represents the log_2_ fold change, and the y-axis shows the –log_10_ p-value. Dots represent individual proteins. Threshold lines indicate significance cutoffs (e.g., fold change > ±0.3 and p < 0.05). Points beyond these thresholds are considered significantly altered. Blue dots represent downregulated proteins, red represent upregulated, and grey represents non-significant. (D) Sankey plot demonstrating shifts in differential protein states in response to hypoxia in the presence or absence of rASPN. Significance categories for each comparison are labeled as upregulated (Log2 fold-change >0.3, FDR <0.05), downregulated (Log2 fold-change <0.3, FDR <0.05), minimal change (Log2 fold-change > −0.3 and <0.3, but FDR <0.05), or non-significant (FDR > 0.05). (E) KEGG pathway enrichment bubble plot across experimental conditions. X axis depicts the pathways divided into four categories: Consensus: Pathways enriched from proteins that were found to be significant and have the same expression profile before and after treatment, Unique: Pathways enriched from proteins that are found to be significant with differentially expressed post rASPN treatment, Reversed: Pathways enriched from proteins enriched from proteins that are found to be significant and expressed prior to treatment but after treatment they are found to change their expression profile, and Suppressed, pathways enriched from proteins which are found to be significant without the treatment are now non-significant on treatment.

We next examined the dynamics of proteome changes in more detail by comparing them across conditions, tracking patterns of protein DA across the treatment groups. To balance practical and statistical significance, we also set a Log2 fold-change (Log2FC) cut-off of >|0.3| to ensure at least modest levels of mean change in a protein, referring to proteins with statistical significance but low overall levels of change as ‘minimally’ affected. The overall results of proteome dynamics are visualized in the Sankey plot in Figure 2D and can be further inspected in Supplementary Table ST1.

In the No-ReOx experiments, among the 1143 proteins significantly altered in H vs N (710 upregulated, 251 downregulated), 35% (n=407, 277 up, 78 down) remained significant in the same direction in the HA vs N comparison. Among these, however, a comparison of HA vs H abundance levels revelated that 29 of the uniformly upregulated or downregulated proteins had a significantly enhanced response (e.g., further upregulated/downregulated in HA vs H beyond the magnitude observed in H vs N) whereas 13 of uniformly altered proteins actually had significantly modulated response in that while they still changed significantly relative to N, the magnitude of the response was less when compared to H alone (e.g., a significantly upregulated protein in HA vs N was significantly downregulated in HA vs H). Along similar lines, just under 42% (n=484) of the hypoxia-response proteins (H vs N, 333 originally up, 151 originally down) were no longer statistically different from non-hypoxic CM when rASPN was added in the presence of hypoxia. A very small number (n=54) demonstrated a significant reversal in their hypoxic response, with 13 upregulated proteins subsequently downregulated with rASPN and 5 downregulated proteins subsequently upregulated. Finally, uniquely significant proteins were identified (n = 200 upregulated, n = 65 downregulated) in the HA vs. N comparison that were not initially altered by hypoxia alone. Taken together, these dynamic patterns of protein change indicate a significant effect of rASPN on modulating the CM response to acute hypoxic stress.

Pathway enrichment analysis of the proteins for each of these patterns was then applied to gain a better understanding of the potential biological implications (Figure 2E). Among the ‘consensus’ hypoxia response proteins (significant in the same direction in both H vs N and HA vs N), several signaling pathways (PI3K-Akt, Rap1), numerous ECM /adhesion-related pathways, as well as lysosomal and mitophagy/autophagy pathways were enriched, pointing towards a core hypoxia response signature that was relatively unchanged by the presence of rASPN. Interestingly, a handful of these same pathways (e.g., ECM, adhesion, autophagy, lysosome) had additional DA proteins only significant in the presence of rASPN (e.g., HA vs N, labelled ‘unique’). This indicates that rASPN may modulate or augment these already enriched functional proteomic responses to hypoxia. Additional pathways enriched only in the HA vs N comparison were included: transforming growth factor beta (TGF-β) and cardiomyopathy. Along with these apparently modulated/augmented and uniquely activated pathways, some proteins involved in several signaling pathways, including Mapk, Rap1, and HIF1a, as well as proteins related to apoptosis, were no longer different relative to normoxic controls in the presence of rASPN (labeled ‘suppressed’). What’s more, we observed some proteins that were not only mollified upon rASPN treatment, but whose direction of DA was reversed (e.g., upregulated in hypoxia alone may be significantly downregulated in hypoxia-rASPN relative to normoxic control). This small set of proteins only showed significant enrichment for the actin-cytoskeletal remodeling pathway.

The analysis reveals that core hypoxia-responsive pathways—such as PI3K-Akt, Rap1, ECM/adhesion, lysosomal, and autophagy processes—remain enriched regardless of rASPN. However, rASPN further enhances or uniquely activates some of these same pathways, including TGF-β and cardiomyopathy signaling. At the same time, rASPN suppresses or reverses the differential abundance of certain proteins involved in pathways like MAPK, Rap1, HIF1α, and apoptosis, with a small subset even showing opposite regulation and enrichment in actin– cytoskeletal remodeling.

### Effect of rASPN on Cardiomyocyte Proteome during Reoxygenation post-Hypoxia

Differential analysis of the ReOx condition identified 902 DA proteins between H and N (12% of the detected proteome, Figure 3A), with a substantially larger proportion of the proteome, 1661 DA proteins (18% of the detected proteome), affected by the addition of rASPN in the hypoxic-recovery interval as compared to the non-hypoxia-exposed control (Figure 3B). Furthermore, rASPN elicited a unique proteomic profile across hypoxia-exposed cells, with 1213 DA proteins (14% of the detected proteome) significantly altered in HA vs H cells (Figure 3C).

**Figure 3.**
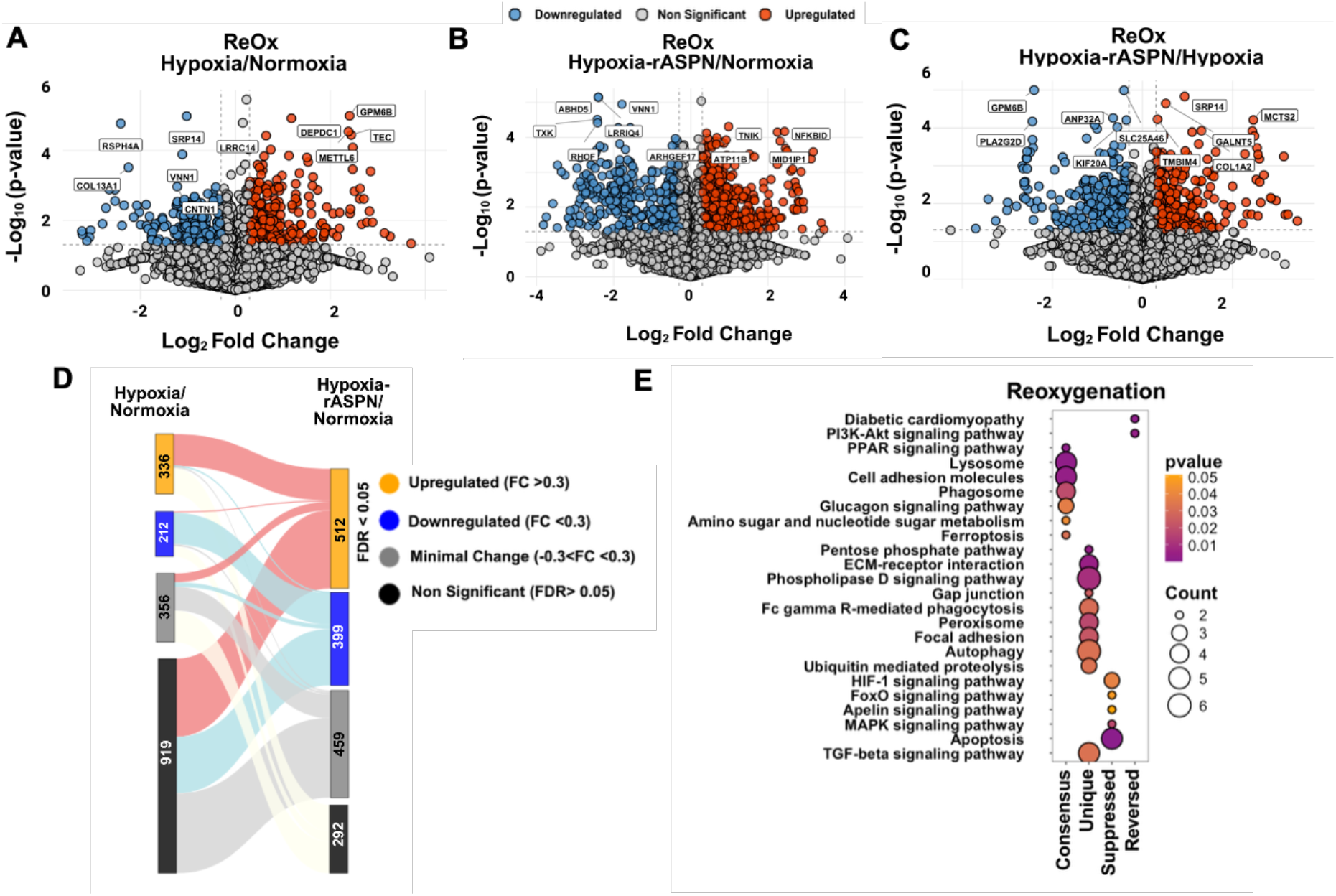
Overview of differentially expressed proteins (DEPs) identified in hypoxia and rASPN-treated conditions under No-ReOx and ReOx. (A) Volcano plots showing differential expression based on (A) the ratio of hypoxia to normoxia for Reoxygenation, (B) the ratio of hypoxia-rASPN to normoxia for Reoxygenation and (C) the ratio of hypoxia-rASPN to hypoxia for Reoxygenation. For volcano plots, the x-axis represents the log_2_ fold change, and the y-axis shows the –log_10_ p-value. Dots represent individual proteins. Threshold lines indicate significance cutoffs (e.g., fold change > ±0.3 and p < 0.05). Points beyond these thresholds are considered significantly altered. Blue dots represent downregulated proteins, red represent upregulated, and grey represents non-significant. (D) Sankey plot demonstrating shifts in differential protein states in response to hypoxia-reoxygenation in the presence or absence of rASPN. Significance categories for each comparison are labeled as upregulated (Log2 fold-change >0.3, FDR <0.05), downregulated (Log2 fold-change <0.3, FDR <0.05), minimal change (Log2 fold-change > −0.3 and <0.3, but FDR <0.05), or non-significant (FDR > 0.05). (E) KEGG pathway enrichment bubble plot across experimental conditions. X-axis denotes protein groups divided into four categories: Consensus: Pathways enriched from proteins that were found to be significant and have the same expression profile before and after treatment, Unique: Pathways enriched from proteins that are found to be significant with differentially expressed post rASPN treatment, Reversed: Pathways enriched from proteins enriched from proteins that are found to be significant and expressed prior to treatment but after treatment they are found to change their expression profile, and Suppressed, pathways enriched from proteins which are found to be significant without the treatment are now non-significant on treatment.

The proteome dynamics analysis was then performed on the ReOx response (Figure 3D, can be further inspected in Supplementary Table ST2), among the 902 proteins significantly altered in H vs N (335 upregulated, 211 downregulated), 50% (n = 451, 136 upregulated, 134 downregulated) remained significant in the same direction in the HA vs N comparison. Among these, however, a comparison of HA vs H abundance levels found 133 of the uniformly upregulated or downregulated proteins had a significantly augmented response (e.g., further upregulated/downregulated in HA vs H relative to response in either H vs N or HA vs N) whereas only 2 of uniformly altered proteins had significantly modulated response in that while they still changed significantly relative to N, the magnitude of the response was less when compared to H alone (e.g., a significantly upregulated protein in HA vs N was significantly downregulated in HA vs H). By contrast, just under 17% (n=156) of these hypoxia-response proteins (111 originally up, 45 originally down) were no longer statistically different from non-hypoxic CM when rASPN was added in the presence of hypoxia. A very small number (n = 56) demonstrated a significant reversal in their hypoxic response, with 37 upregulated proteins subsequently downregulated in response to rASPN and 19 downregulated proteins subsequently upregulated. We also identified uniquely significant proteins (n = 334 upregulated, n = 243 downregulated) in the HA vs. N comparison that were not initially altered by hypoxia alone. Taken together, these dynamic patterns of protein change indicate a significant effect of rASPN on modulating the CM response to hypoxia-reoxygenation stress.

As described above for acute hypoxia (e.g., NoReOx); pathway analysis was used to infer possible functional implications of these proteomic patterns of change (Figure 3E). Among the ‘consensus’ hypoxia response proteins (significant in the same direction in both H vs N and HA vs N), several signaling pathways (PPAR, Rap1), numerous ECM /adhesion-related pathways, as well as lysosomal and mitophagy/autophagy pathways were enriched, pointing towards a core hypoxia-reoxegenation response that was relatively unchanged by the presence of rASPN. Interestingly, a handful of these same pathways (e.g., ECM, adhesion, autophagy, lysosome, Focal adhesion, Peroxisomes, and Gap junction) had additional DA proteins only significant in the presence of rASPN (e.g., HA vs N, labelled ‘unique’). This indicates that rASPN may modulate or augment these already enriched functional proteomic responses to hypoxia-reoxygenation. Additional pathways enriched only in the HA vs N comparison were included: transforming growth factor beta (TGF-β). Along with these apparently modulated/augmented and uniquely activated pathways, some proteins involved in several signaling pathways, including Mapk, FoxO, and HIF1a, as well as proteins related to apoptosis, were no longer different relative to normoxic controls in the presence of rASPN (labeled ‘suppressed’). What’s more, we observed some proteins that were not only mollified upon rASPN treatment, but whose direction of DA was reversed (e.g., those upregulated in hypoxia alone may be significantly downregulated in hypoxia-rASPN relative to the normoxic control). This small set of proteins showed significant enrichment only for the PI3K-Akt and diabetic cardiomyopathy signaling pathways.

Overall, the proteomic analysis suggests that the addition of rASPN during the reoxygenation phase strengthens metabolic, autophagic, and stress-response pathways, and rASPN further enhances ECM/adhesion and autophagy pathways while lowering the enrichment of MAPK, Rap1, Hif-1alpha, apoptosis, and shifting the cytoskeletal stress signaling, indicating a selective modulation of CMs recovery during the reoxygenation phase. Taken together, this pattern of functional pathway enrichment informs our hypothesis that rASPN promotes CM survival after hypoxia-reoxygenation, at least in part, by augmenting autophagy and subsequently inhibiting apoptosis and/or promoting cell survival signaling. We next set out to validate these proteomic findings and further test this hypothesis. Given that rASPN treatment is likely most practical to administer during the reperfusion (e.g., reoxygenation) interval in a human patient, we focused our validation experiments on the ReOx *in vitro* model.

### Validation of proteomic-predicted effects of rASPN on CM autophagy and apoptosis *in vitro* during ischemic reperfusion

Autophagy pathway proteins were identified in our proteomics data in both the NoReOx and ReOx conditions. Given our previous work highlighting the role of rASPN in regulating autophagy in cardiac fibroblasts, as well as the well-established role of autophagy in modulating apoptosis, we sought to further investigate these two pathways in this hypoxic CM model.

First, we tested whether the upregulation of autophagy-related proteins was indicative of actual changes in autophagic flux. Importantly, the key autophagic regulator, BNIP3, was detected as a rASPN-modulated autophagy member in our proteomic analysis (Figure 4A). Western blotting confirmed and extended these findings (Figure 4B), demonstrating a strong upregulation of both BNIP3 and Beclin-1 in CMs treated with rASPN during the reoxygenation interval.

**Figure 4.**
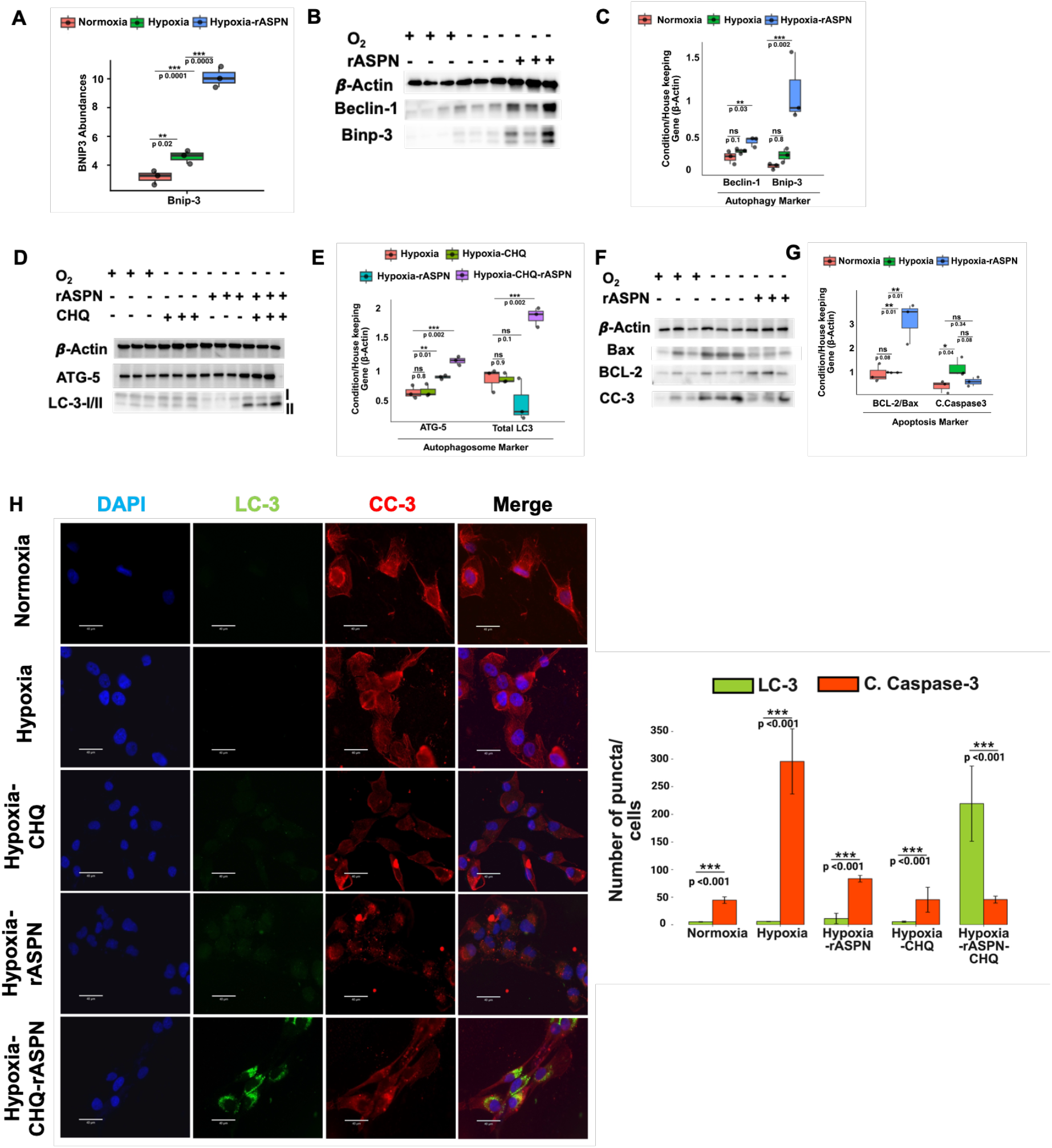
Characterization of novel protein analysis under hypoxia and treatment with exogenous rASPN conditions.(A)Protein abundances of HIF-1α across all proteomics data analysis across Normoxia, Hypoxia, and Hypoxia-rASPN for ReOx (p-value presented on the graph). (B) Western blot analysis showing increased expression of β-Actin, Beclin-1, BNIP3, and LC3-I/II levels; Normoxia, Hypoxia, and Hypoxia-rASPN (p-value presented on the graph). (C) Densitometry quantification showing increased expression of β-Actin, Beclin-1, BNIP3, and LC3-I/II levels; Normoxia, Hypoxia, and Hypoxia-rASPN, of blots in Figure 3B. (D) Western blot data showing increased LC3-II levels in the presence of chloroquine (CHQ), with elevated expression of autophagosome-related genes ATG5 and ATG7 upon rASPN treatment (p-value presented on the graph). (E) Densitometry quantification showing increased expression of LC3-II, upon rASPN treatment in the presence of CHQ. (F) Western blot analysis reveals increased Actin and BCL-2 levels, and decreased BAX levels, as well as CC-3, indicating suppression of apoptosis and an increase in autophagy following rASPN treatment (G) Densitometry quantification revealed increased BCL-2: BAX ratio levels and CC-3, indicating suppression of apoptosis and an increase in autophagy following rASPN treatment (p-value presented on the graph). (H) Immunofluorescence (IF) analysis of cells under Normoxia, Hypoxia, Hypoxia-rASPN, Hypoxia in the presence of CHQ, and Hypoxia-rASPN in the presence of CHQ, highlighting the localization and expression dynamics of autophagy/apoptosis markers (p-value presented on the graph).

To determine whether autophagic flux is indeed upregulated by rASPN, we examined levels of the autophagosome-formation markers ATG5 and LC-3-II in the presence (or absence) of lysosomal inhibitor, chloroquine (CHQ), which blocks the final degradation of autophagosomes and thus permits accumulation of these key readout markers even under conditions of high flux. As shown in Figures 4D and 4E, while we saw little indication of functional autophagy in the ReOx-treated CMs alone, the addition of rASPN during reoxygenation slightly increased the early autophagy marker Atg5, an effect that was amplified upon inhibition of degradation by CHQ. Interestingly, the functional marker LC3 was markedly absent from detection in rASPN-only-treated ReOx CMs, but was profoundly abundant after the addition of CHQ, indicating increased autophagic flux and lysosomal degradation in the presence of rASPN that became detectable upon inhibition of final degradation by CHQ.

Since autophagy is a well-known regulator of CM apoptosis, and apoptosis pathways were also observed to be influenced by rASPN in our proteomics analysis, we then examined levels of the apoptosis signaling indicators BCL2, Bax, and Cleaved Caspase 3 (CC3) by western blot (Figure 4F and Figure 4G). Consistent with an increase in apoptotic signaling, CMs exposed to hypoxia and then reoxygenation also significantly upregulated both BAX and CC3, which was significantly blunted by the addition of rASPN during the reoxygenation interval. Interestingly, we observed an increase in BCL2 expression in ReOx CMs, with further and significant elevation by rASPN. With these patterns, the combined BCL2/BAX ratio was consistent with elevated apoptosis signaling provoked by hypoxia-reoxygenation, which was suppressed by the addition of rASPN (Figure 4G).

To further confirm the role of rASPN in modulating the autophagy and apoptosis responses of AC16 CMs to hypoxia-reoxygenation, we assessed the spatial accumulation of CC3 and LC3 using immunofluorescence (IF). We quantified the accumulation of CC3 and LC3 into perinuclear puncta, as indicative of active apoptosis signaling and autophagy, respectively (Figure 4H). Accordingly, we observed a higher number of LC-3 puncta and fewer CC3 puncta near the nucleus in the case of Hypoxic cells treated with rASPN in the presence of CHQ compared to other conditions (Normoxia, hypoxia, hypoxia-CHQ, and hypoxia-rASPN).

### ASPN is associated with altered apoptosis and autophagy in a murine myocardial infarction model *in vivo*

To confirm the relevance of these effects observed *in vitro* to an *in vivo* cardiac paradigm, we leveraged existing tissue work related to our previous publication [15] describing mice with normal (WT) and genetically absent ASPN (ASPN-KO) that had been subjected to permanent coronary artery ligation (PCAL)-induced heart damage. Immunohistochemistry-fluorescence (IHC-F) was used to detect the markers for autophagy (LC-3 (Green fluorophore)) and apoptosis (CC-3(Red fluorophore)) (Figure 5A). We computationally identified cells based on DAPI-positive nuclei and quantified the puncta of each marker (Figure 5B), comparing the sham to both direct infarct and infarct-remote regions of the PCAL-treated mice. We observed that while WT mice exhibit an upregulation of LC3 in the infarct-damaged myocardium, the genetic absence of ASPN completely blocks this response (Figure 5B, upper right panel). Interestingly, minimal LC3 puncta were observed in sham or remote regions, though small but significant differences were still observed between WT and ASPN-KO. In contrast to autophagy, genetic loss of ASPN promoted a stronger CC-3 signal across all conditions relative to WT, with nearly twice the increase in CC-3 signal in the infarcted region compared to the sham or remote region. Thus, whereas our *in vitro* data show that adding ASPN to CMs promotes autophagy and inhibits apoptosis in response to hypoxic conditions, these *in vivo* data indicate that loss of ASPN demonstrates the reverse, with lower apparent autophagy and higher apoptosis signaling, especially in the direct infarct region, presumably most impacted by hypoxia exposure.

**Figure 5.**
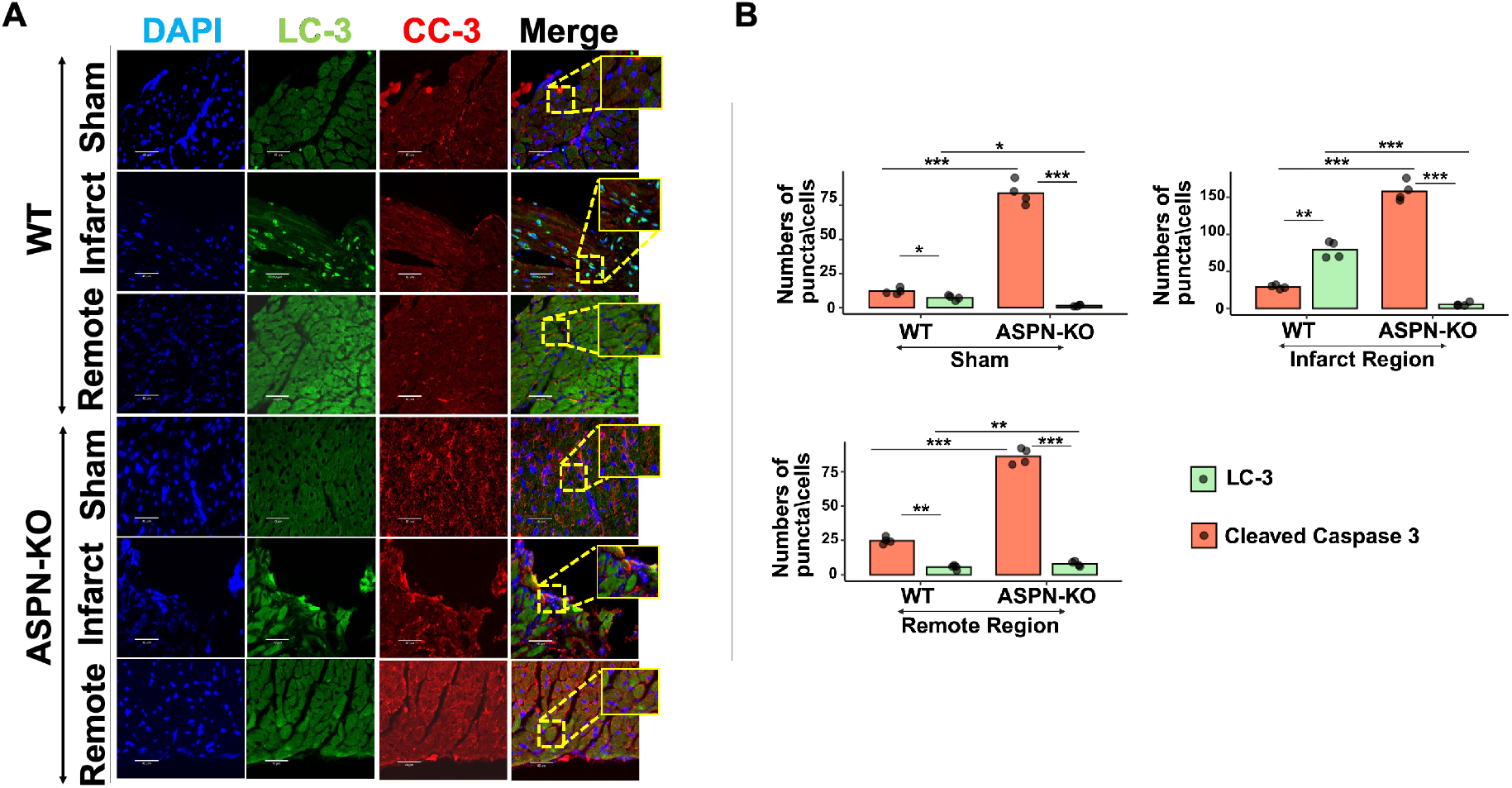
Immunohistochemistry-fluorescence (IHC-F) analysis in PCAL mouse models for both WT and ASPN-KO mice. (A) Immunohistochemistry-fluorescence (IHC-F) analysis of heart samples of both sham and PCAL for WT and ASPN-KO mice for 24 hours. In the case of Ischemic-reperfusion injury (IRI), there are two distinct zones: the infarct region and remote regions. highlighting the localization and expression dynamics of autophagy/apoptosis markers. Further puncta near the nucleus and around the cytoplasm for both the markers were estimated and plotted. (B) Puncta analysis for LC-3 and CC-3 for both wild-type sham (WT-Sham), ASPN knockout sham (KO-Sham), wild-type PCAL (WT-PCAL), and ASPN knockout PCAL (KO-PCAL), presented as the sham group, infarct region group, and Remote region group.

### rASPN treatment reverses the TGF-β pathway in both No-ReOx and ReOx ischemic reperfusion *in vitro* model

In addition to the autophagy/apoptosis effects, functional pathway analysis on proteomics data indicated a unique enrichment of TGF-β signaling related proteins among those uniquely significant in both acute and ReOx HA vs N comparisons.

We used Western blotting to determine how rASPN affects key TGFβ signaling indicators, TGFB1 ligand, and the signaling effectors SMAD2/3 (Figure 6A & 6B). Western blot analysis for TGF-β1 ligand did not indicate differences across groups; however, levels of SMAD2/3 phosphorylation were elevated under ReOx and further potentiated in the presence of rASPN. Upon phosphorylation by TGF-βR, SMAD2 and/or 3 dimerize with SMAD4 and localizes to the nucleus to modulate gene expression [24]. To examine whether the observed patterns of SMAD2/3 phosphorylation are consistent with changes in nuclear localization of SMAD4 and, by inference, active TGFβ-mediated gene regulation, we assessed the nuclear versus cytosolic localization of SMAD4 using IF (Figure 6C). Interestingly, while SMAD2/3 phosphorylation was increased in both ReOx alone and further amplified by rASPN, we observed prominent SMAD4 nuclear localization only in the ReOx condition without rASPN, as quantified by the nuclear-to-cytosolic ratio of SMAD4 signal intensity (Figure 6D), indicating that rASPN blocks nuclear localization under these conditions.

**Figure 6.**
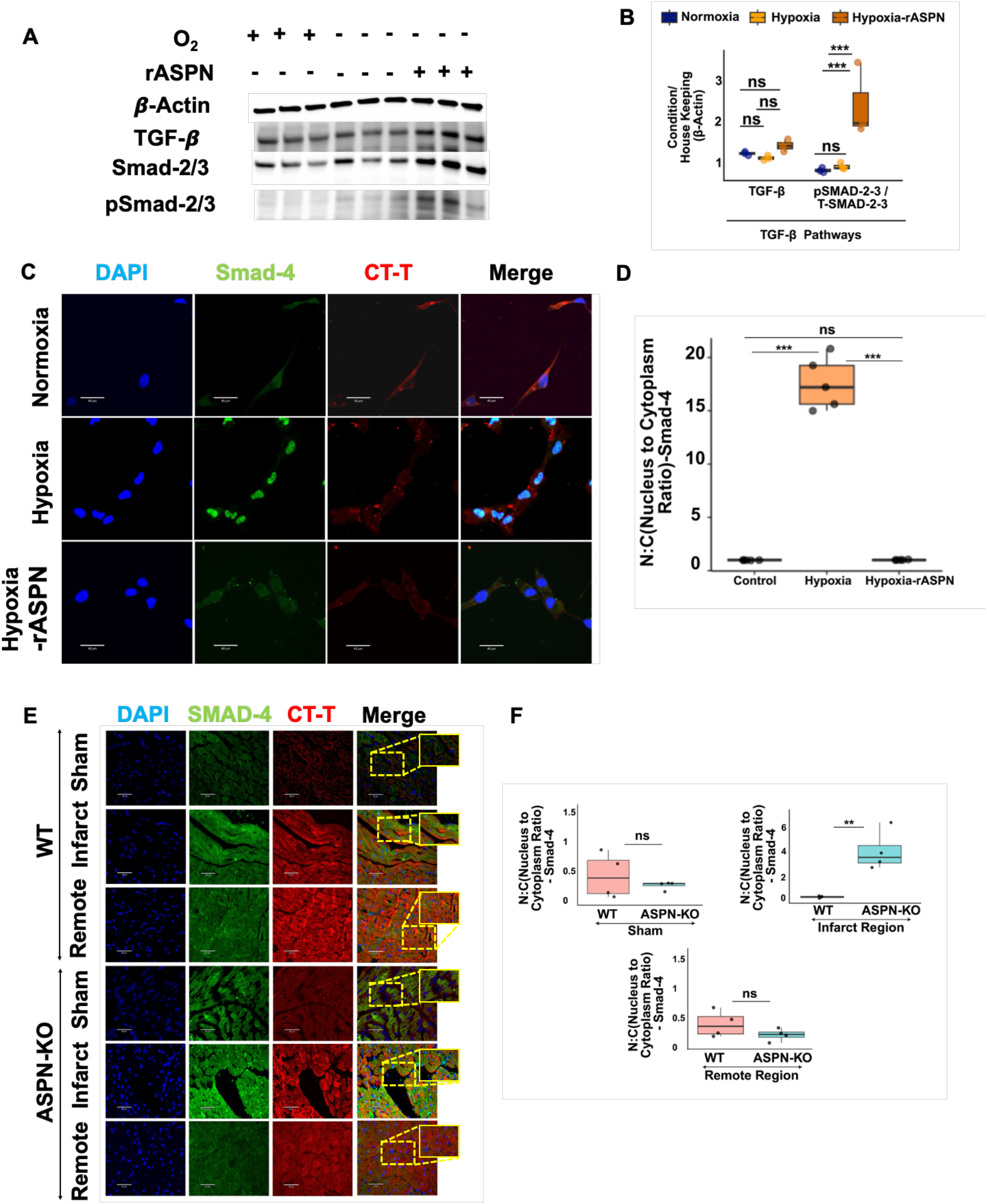
Western blot and densitometry analysis of protein from TGFβ pathways, which were part of novel pathways. (A) Western blot analysis showing increased expression of β-Actin, TGF-βR1, Smad-2/3, and Psmad-2/3 levels; Normoxia, Hypoxia, and Hypoxia-rASPN. (B) Densitometry analysis of TGF-βR1, and ratio of total-Smad-2/3 and Psmad-2/3 levels expression in all three conditions, and normalized with β-Actin (Housekeeping). (C) Immunofluorescence (IF) analysis of cells under Normoxia, Hypoxia, hypoxia-rASPN, highlighting the localization and expression of SMAD-4 in the nucleus and cytoplasm, estimation of nuclear and cytoplasm localization of Smad-4, and further calculation of the ratio of them for each condition.(D) Puncta analysis for SMAD-4 in cells under Normoxia, Hypoxia, hypoxia-rASPN, highlighting the localization and expression of SMAD-4 in the nucleus and cytoplasm, estimation of nuclear and cytoplasm localization of Smad-4, and further calculation of the ratio of them for each condition.(E) Immunohistochemistry-fluorescence (IHC-F) analysis of heart samples of WT and ASPN-KO mice in two different zones near the infarct region and remote regions for both mice under Sham and Ischemic-reperfusion injury (PCAL). Localization and expression of SMAD-4 in the nucleus and cytoplasm, estimation of nuclear and cytoplasm localization of Smad-4, and further calculation of the ratio of them for each condition.(F)Puncta analysis for SMAD-4 for both wild-type sham (WT-Sham), ASPN knockout sham (KO-Sham), wild-type PCAL (WT-PCAL), and ASPN knockout PCAL (KO-PCAL), presented as the sham group, infarct region group, and Remote region group.

To extend this intriguing potential regulatory effect of ASPN on TGFB to an *in vivo* system, we assessed SMAD4 expression in the PCAL mouse model described above, comparing WT and ASPN-KO mice across Sham, Infarct-near, and Infarct-remote regions (Figure 6E). Cardiac Troponin T staining was used to ensure only CMs were assessed for SMAD4 signal. In the absence of ASPN, we observed higher SMAD4 localization in the nucleus of cells within the infarct region compared to WT mice (Figure 6E). There was no obvious increase in SMAD4 localization in the sham or remote regions, as quantified by the nuclear-to-cytosolic ratio of SMAD4 signal intensity (Figure 6F).

## Discussion

In this study, we hypothesized that rASPN, a recombinant peptide that mimics the ECM protein ASPN, plays a significant role in modulating CM responses to acute hypoxia and ischemia-reperfusion (I/R). To test this, we utilized an *in vitro* model to test the acute hypoxia response after 18 hours and the ischemia/reperfusion response after both 18 hours of hypoxia and 12 hours of normoxic recovery. Through in-depth proteomic profiling, we identified numerous functional pathways in both the acute hypoxia and ischemia-reperfusion models that were modulated by the addition of rASPN. Among these, we highlighted and confirmed the importance of rASPN treatment in promoting autophagy and mitigating apoptotic signaling, thereby confirming our previous report on the role of ASPN in regulating these pathways and extending our earlier findings in fibroblasts to demonstrate additional pro-survival benefits directly in CMs.

Autophagy plays a protective role during ischemic stress in cardiac cells by removing damaged organelles and supporting energy metabolism [25]. Our data demonstrate that rASPN enhances BNIP-3-mediated autophagy in CMs. As evidenced by the increased upregulation of BNIP-3 and Beclin-1, as well as increased CHQ-dependent LC3-II accumulation, the direct mechanism through which this occurs remains unknown. Through past research, it has been observed that BNIP-3 overexpression results in the substantial removal of mitochondria by autophagosomes in adult cardiac myocytes, thereby inducing autophagy [26-28]. However, in this study, we have not investigated the impact of rASPN on the ubiquitin-mediated pathway; instead, we have focused on experimental conditions that can differentiate between global autophagy and mitophagy.

Autophagy has been closely linked to the regulation of apoptosis in cardiomyocytes and other cell types [29, 30]. Consistent with these prior observations, we observed that rASPN inhibits apoptosis, as evidenced by reduced BAX expression, increased BCL-2 expression, and decreased cleaved caspase-3 signals, as determined by western blot analysis. These contrasting trends observed between autophagy and apoptosis in CMs on treatment with rASPN suggest to us that ASPN acts as a protective agent in CMs during cardiac stress. The autophagy-apoptosis balance is crucial for CM survival during ischemia-reperfusion (I/R) injury. While we did not experimentally determine whether rASPN inhibits apoptosis in the absence of its effects on autophagy, our findings are consistent with ASPN having a role as a central modulator of this balance and warrant further testing in myocardial ischemia-reperfusion models.

Another important pathway that we observed to be modulated by rASPN was the TGF-β signaling pathway. Our data showed no impact of ASPN on the TGF-β ligand; however, intriguingly, despite upregulating SMAD2/3 phosphorylation, ASPN also apparently blocked nuclear SMAD complex localization. This is an intriguing mechanistic paradox, indicating complex regulatory capacity for ASPN. One possible explanation for reduced nuclear localization despite increased SMAD activation could be the upregulated autophagic state of the ASPN-treated CMs. SMAD4 or the SMAD2/3-SMAD4 complex may be targeted (e.g., through the p62 pathway) for sequestration and lysosomal degradation. The potentiation of CM SMAD2/3 phosphorylation directly contradicts our previous report, which showed augmented SMAD2 activation in cardiac fibroblasts with ASPN knockdown [15]. The differences between this previous finding and our current result may indicate cell-type specificity in how ASPN modulates TGF-β within a tissue or may also be explained by technical factors such as the hypoxic pre-treatment of the CMs (versus non-hypoxic fibroblasts previously), or even the species of cell used (human here versus mouse previously). Interestingly, the net outcome of reduced TGF-β signaling is consistently observed across studies, as the nuclear localization and subsequent transcriptional activation of TGF-β response genes were blocked by ASPN in our CMs. The duality in TGF-β modulation that we observed in our experiments may be consistent with the complex role of this signaling pathway in myocyte ischemia-reperfusion response, with both beneficial and detrimental effects observed on overall outcomes [31]. Our data suggest that ASPN plays a complex modulatory role, and depending on other contextual factors, can either augment or suppress TGF-β signaling and transcriptional activity. The implications of this warrant further investigation in future studies.

We also observed that rASPN has the ability not only to modulate proteins previously affected by hypoxia but also to induce significant changes in proteins that were not initially impacted. This emphasizes that rASPN could rewire the CM’s response beyond hypoxia-induced damage. In line with this, we observed that in both ischemic models, under hypoxic conditions, we observed that rASPN uniquely impacts key signaling cascades associated with cytoskeletal remodeling (Regulation of the actin cytoskeleton), metabolic adaptation (Valine-leucine-isoleucine degradation), ECM remodeling (ECM receptor interaction), and DNA damage responses (p53 signaling) relative to hypoxia alone. Taken together with our findings related to autophagy, apoptosis, and TGF-β, these results further highlight ASPN’s role in modulating cellular homeostasis in CMs within the context of hypoxia and reoxygenation. While we did not focus on these additional functions in the current work, these may become additional areas of future study into how this and other matrix proteins modulate the CM stress response.

As a preliminary extension of our findings to the *in vivo* paradigm, we examined whether the beneficial effects of adding rASPN to CMs observed *in vitro* would show opposite trends *in vivo in mouse* hearts with and without genetic ASPN loss during cardiac ischemia induced by PCAL. Indeed, this was the case for all three of the major effects examined; LC3 appeared higher and CC3 levels lower in the infarcted region of WT heart relative to ASPN KO, and while *in vitro* SMAD4 nuclear localization was absent with rASPN treatment, *in vivo* we observed significantly higher levels of nuclear SMAD4 in ASPN-KO infarct cells relative to WT. We also know from our previous report that ASPN-KO mice have significantly impaired functional recovery, with exacerbated fibrosis and larger infarct regions, after PCAL relative to WT mice [15]. Our current analysis extends these findings to indicate the direct, *in situ* relevance of autophagy, apoptosis, and TGF-β signaling to these previously reported functional observations, and extends them to elucidate key cardiomyocyte specific effects, observed from our *in vitro* experiments. The provisional *in vivo* results reported here, coupled with our strong *in vitro* effects, provide a strong rationale to further interrogate rASPN as a therapeutic agent in future studies.

## Conclusion

Overall, these findings suggest that rASPN exerts a multifaceted protective role in CMs. CMs, when treated with rASPN under hypoxic conditions, modulate a vast network of signaling pathways that involve the modulation of several key pathways, including BNIP-3-mediated autophagy, apoptosis, metabolism, and CM cytoskeletal/ECM remodeling. While we have yet to trace the direct receptor for and downstream effects of rASPN, our results provide a strong rationale for further investigation of rASPN as a potential therapeutic agent for myocardial ischemia and related pathologies. Importantly, our ReOx model highlights ASPN’s therapeutic potential during reperfusion, where it could be added during vascular interventions to clear blockages (e.g., coronary artery bypass grafts, coronary artery stenting) to enhance CM and tissue healing in the reperfusion phase, positioning it as a promising candidate for future cardioprotective interventions in ischemic heart disease.

## Authorship contribution statement

Conceptualization of experimental design, data analysis, and interpretation was done by DR and SJP with early support from HP. Supervision is done by SJP and ABS. Sample collection is done by DR and MB. Proteomics sample processing is done by DR, LM, MA, and AB. DR, AD, and DG performed IF experiments and analyzed the data. PN helped with slide preparation for IF.

## Declaration of competing interests, generative AI, and AI-assisted technologies in the writing process

The authors declare that they have no known competing financial interests or personal relationships that could have appeared to influence the work reported in this manuscript. The authors have not used AI-assisted techniques in this manuscript.

## Ethical Approval

All animal experiments were designed and performed in compliance with the National Institutes of Health and were approved by the Institutional Animal Care and Use Committee of Cedars-Sinai Medical Center under the protocol IACUC-008858.

## Acknowledgements

We would like to acknowledge the Proteomics Core, Cedars-Sinai Biobank, and Translational Research Core for histology services. We thank Lior Zilberberg for his valuable input during the data discussion and Ian Williamson for his assistance in excellent lab management. We thank Roberta A. Gottlieb, Honit Piplani, Reetu Thankur, and Chengqun Huang for their previous work on the PCAL model.

